# Bacterial ADP-heptose initiates a revival stem cell program in the intestinal epithelium

**DOI:** 10.1101/2024.01.15.575749

**Authors:** Shawn Goyal, Cynthia X. Guo, Adrienne Ranger, Derek K. Tsang, Ojas Singh, Caitlin F. Harrigan, Olga Zaslaver, Hannes L. Rost, Herbert Y. Gaisano, Scott A. Yuzwa, Nan Gao, Jeffrey L. Wrana, Dana J. Philpott, Scott D. Gray-Owen, Stephen E. Girardin

## Abstract

The intestinal epithelium has an exceptional capacity to repair following injury, and recent evidence has suggested that YAP-dependent signaling was crucial for the expansion of *Clu*^+^ revival stem cells (revSCs) with fetal-like characteristics, which are essential for epithelial regeneration. However, neither the mechanism underlying where these revSCs emerge from nor the nature of the physiological cues that induce this revSC program, are clearly identified. Here, we first demonstrate that *Alpk1* and *Tifa*, which encode the proteins essential for the detection of the bacterial metabolite ADP-heptose (ADP-Hep), were expressed by the stem cell pool in the intestinal epithelium. Treatment of intestinal organoids with ADP-Hep not only induced acute NF-κB pro-inflammatory signaling but also TNF-dependent apoptosis within the crypt, causing blunted proliferation and acute disruption of the crypt architecture, while also triggering induction of a revSC program. To identify the molecular underpinnings of this process, we performed single-cell RNA-seq analysis of ADP-Hep-treated organoids as well as lineage-tracing experiments. Our data reveal that ADP-Hep induced the specific ablation of the homeostatic intestinal stem cell (ISC) pool. Removal of ADP-Hep resulted in the rapid recovery of ISCs through dedifferentiation of Paneth cells, which transiently acquired revSC features and expressed nuclear YAP. Moreover, lineage tracing from *Lyz1*^+^ Paneth cells showed that ADP-Hep triggered Paneth cell de-differentiation towards pluripotent and proliferative cells in organoids. *In vivo*, revSC emergence in response to irradiation-induced injury was severely blunted in *Tifa*-deficient mice, suggesting that efficient epithelial regeneration in this model required detection of microbiota-derived ADP-Hep by the ALPK1-TIFA pathway. Together, our work reveals that Paneth cells can serve as the cell of origin for revSC induction in the physiological context of microbial stimulation, and that the transient loss of *Alpk1*-expressing ISCs is the initiating event for this regenerative process.

---

A monolayer of epithelial cells lines the intestinal tract and provides an essential physical barrier shielding the mucosa from the intestinal lumen where the microbiota resides. Intestinal epithelial cells (IECs) also secrete factors, such as mucins (produced by goblet cells) and a wide variety of antimicrobial peptides and proteins (produced by Paneth and goblet cells, mainly) that further reinforce the epithelial barrier. To maintain the integrity of this epithelial barrier, IECs are very rapidly turned over (in approx. 3-5 days), a consequence of the intense proliferative capacity of intestinal stem cells (ISCs). ISCs are thus essential elements of the intestinal epithelial barrier and, as such, protecting these cells from enteric pathogens or other insults is paramount ^1^. This is in part achieved by the topology of the intestinal crypt: ISCs are found at the bottom of these crypts, intercalated between Paneth cells, in an area that forms a niche that remains essentially sterile ^2^. At homeostasis, ISCs proliferate actively to give rise to secretory (Paneth, goblet, enteroendocrine, tuft) cells and absorptive enterocytes, and express cellular markers, such as *Lgr5* ^3^.

How ISCs respond to epithelial injury is an area of intense investigation, as this question is crucial for understanding the pathophysiology of enteric infections or chronic inflammatory bowel disease (IBD), for which epithelial injury and barrier breach are key common features ^4^. Recent studies demonstrated that, following acute intestinal injury induced by whole body γ-irradiation (a common model used to study IEC injury and restitution), a specific type of ISCs rapidly emerged through activation of the YAP/TAZ signaling pathway ^5,6^. These ISCs were coined as “revival stem cells” or revSCs, express cellular markers including *Clu*, and are required for the restitution of the intestinal epithelium after injury. Other studies have identified that epithelial injury resulted in the emergence of fetal-like ISCs, as they were found to express cellular markers typically observed in the embryonic intestine, including *Ly6a* ^7,8^. It is highly probable that revSCs and fetal-like ISCs represent either the same or overlapping cell populations; indeed, both are induced by injury and are dependent on YAP signaling. Moreover, revSCs induced following irradiation express both *Clu* and *Ly6a* ^6^. We will thus refer to revSCs/fetal-like ISCs as “revSCs” from now on for simplicity. However, the mechanisms underlying revSC generation and the physiological triggers that control the emergence of these cells remain incompletely characterized. Specifically, the contribution of the microbiome and of microbiota-associated metabolites to revSC induction has not been identified.

Here we identified that the bacterial metabolite ADP-Hep, an intermediate of Gram-negative bacterial lipopolysaccharide (LPS) biosynthesis, which is detected by the host through the ALPK1-TIFA pathway ^9,10^, triggers the acute loss of the ISC pool through apoptosis, thereby driving the induction of the revSC program in IECs. Furthermore, we demonstrate that revSC induction following ADP-Hep treatment occurs through the de-differentiation of Paneth cells. *In vivo*, we found that ADP-Hep was constitutively produced by the intestinal microbiota and that revSC emergence following γ-irradiation was dependent on the ALPK1-TIFA pathway.

## The Alpk1-Tifa signalling pathway is expressed and functional within the intestinal epithelium

We first mined a published RNAseq dataset (GSE81125 ^11^) that compared gene expression in the small intestinal (SI) tissue between conventionally-raised and antibiotic-treated mice, to identify candidate microbial sensors that could modulate the activity of the ISC pool upon colonization with the intestinal microbiota. Interestingly, we noticed that *Alpk1* and *Tifa* were strikingly upregulated by microbial colonization (Fig. 1a). In agreement, a second dataset (GSE60163 ^12^) comparing germ-free (GF) and conventionally-raised mice also revealed significant induction of these genes by the microbiota (Fig. 1b). By analyzing the scRNAseq dataset of isolated intestinal crypts that we recently published ^13^, we further observed that both *Tifa* and *Alpk1* were expressed in a discrete subset of ISCs that shows high expression of the proliferation marker *Mki67* and of the ISC marker *Lgr5* (Fig. 1c). Dual RNA *in situ* hybridization (RNA-ISH) for *Tifa* and *Alpk1* confirmed co-expression with *Lgr5* in intestinal crypts (Fig. 1d).

**Figure 1.**
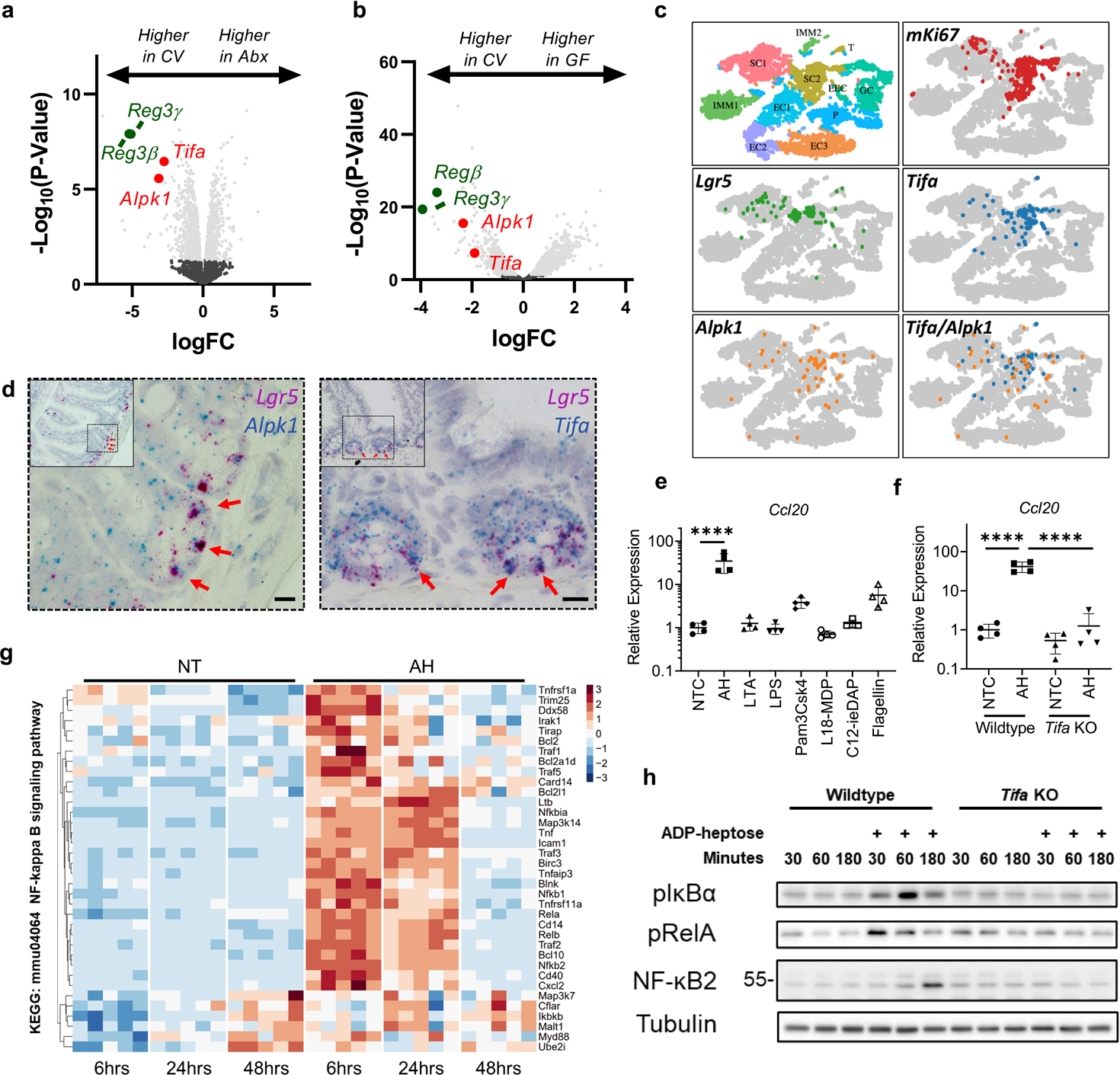
The ALPK1-TIFA axis is expressed and functional within ISCs in the small intestine. **a,** Volcano plot of differential mRNA expression of jejunal tissue from litter mate mice that were either conventionally raised or administered broad spectrum antibiotic in drinking water. Analysis is based on raw data from NCBI GEO: GSE81125. **b,** Volcano plot of differential mRNA expression of small intestinal tissue from either conventionally raised or germ-free mice. Differential expression data is from NCBI GEO: GSE60163. **c,** UMAP from single-cell RNA sequence data of mouse intestinal epithelial cells. **d,** Representative images from dual RNA-*in situ* hybridization (RNA-ISH) for *Lgr5* in red or *Tifa/Alpk1* in blue. Red arrows indicate areas of co-localization (n = 3 mice). **e,** RT-qPCR of ileal organoids stimulated with MAMPs at 10µg/mL for 2 hours (n = 4). **f,** RT-qPCR of ileal organoid generated from either wildtype or *Tifa^-/-^* mice, treated with ADP-Hep for 2 hours (n = 4). **g,** Heatmap generated from bulk RNA-seq data of wildtype ileal organoids untreated or treated with ADP-Hep for 6, 24 or 48 hours. Transcripts are from the KEGG term: NF-kappa B signaling pathway. **h,** Representative immunoblot of ileal organoids from either wildtype or *Tifa^-/-^* mice, stimulated with ADP-Hep for 30, 60 or 180 minutes (n = 3). ****P<0.0001. Scale bar 10µm (d).

We next investigated whether the ALPK1-TIFA pathway was functional in IECs. Intestinal organoids were stimulated with an array of microbe-associated molecular patterns (MAMPs) at equal concentrations, including ADP-Hep, the bacterial ligand of the ALPK1-TIFA pathway, as well as lipoteichoic acid (LTA; TLR2 agonist), LPS (TLR4 agonist), Pam3CSK4 (TLR1/2 agonist), L18-muramyl dipeptide (L18-MDP; NOD2 agonist), c12-iE-DAP (NOD1 agonist) and flagellin (TLR5 agonist). While LTA, LPS, L18-MDP and c12-iE-DAP did not upregulate the expression of the proinflammatory gene *Ccl20* and Pam3CSK4 and flagellin induced moderate *Ccl20* expression, ADP-Hep was by far the most potent trigger (Fig. 1e). As expected, ADP-Hep-dependent induction of *Ccl20* was fully abrogated in organoids from *Tifa^-/-^* mice (Fig. 1f).

To gain a broader understanding of the effects ADP-Hep on the intestinal epithelium, we next performed bulk RNAseq on wild type (WT) organoids treated with ADP-Hep for 6, 24 and 48 hours. At the 6-hour time point, ADP-Hep induced a robust NF-κB-associated proinflammatory gene signature (KEGG: mmu04064) that persisted at the 24-hour time point but was reduced at 48 hours (Fig. 1g and Supplementary Fig. 1a). In agreement, immunoblot analysis of organoids from WT and *Tifa*^-/-^ mice stimulated with ADP-Hep demonstrated rapid induction of phospho-IκBα (pIκBα) and phospho-RelA (pRelA) at 30 minutes of treatment, and of p52 NF-κB2 at 180 minutes, in a TIFA-dependent manner (Fig. 1h). Together, these data demonstrate that the ALPK1-TIFA signalling axis is expressed within ISCs and that stimulation of this pathway triggers NF-κB-associated induction of a pro-inflammatory gene expression program.

## ADP-Hep inhibits epithelial proliferation in intestinal organoids

Our bulk RNAseq dataset revealed that, in addition to the upregulation of NF-κB-dependent transcription, expression of genes associated with DNA replication (KEGG: mmu03030) was severely blunted by ADP-Hep treatment (Fig. 2a; Supplementary Fig. 1b), suggesting that this bacterial molecule might also affect cell proliferation. In support, ADP-Hep significantly reduced 5-ethynyl-2’-deoxyuridine (EdU) incorporation in WT organoids (Fig. 2b). Moreover, ADP-Hep treatment caused a dose-dependent blunting of *Lgr5* expression, suggesting a direct impact on the ISC pool (Fig. 2c). To assess the functional effect of ADP-Hep on IEC growth, we used brightfield microscopy to visualize intestinal organoids at 6, 24 and 48 hours of treatment. While untreated WT organoids exhibited growth associated with the formation of multiple crypts over time, ADP-Hep-treated organoids were severely atrophied, and this effect was TIFA-dependent since *Tifa^-/-^* organoids treated with ADP-Hep grew normally (Fig. 2d, e). In support, ADP-Hep treatment resulted in a marked decrease in Ki67 staining, as visualised by immunofluorescence, in WT but not *Tifa^-/-^* organoids (Fig. 2f). Overall, these data reveal that ADP-Hep potently suppresses intestinal epithelial proliferation.

**Figure 2.**
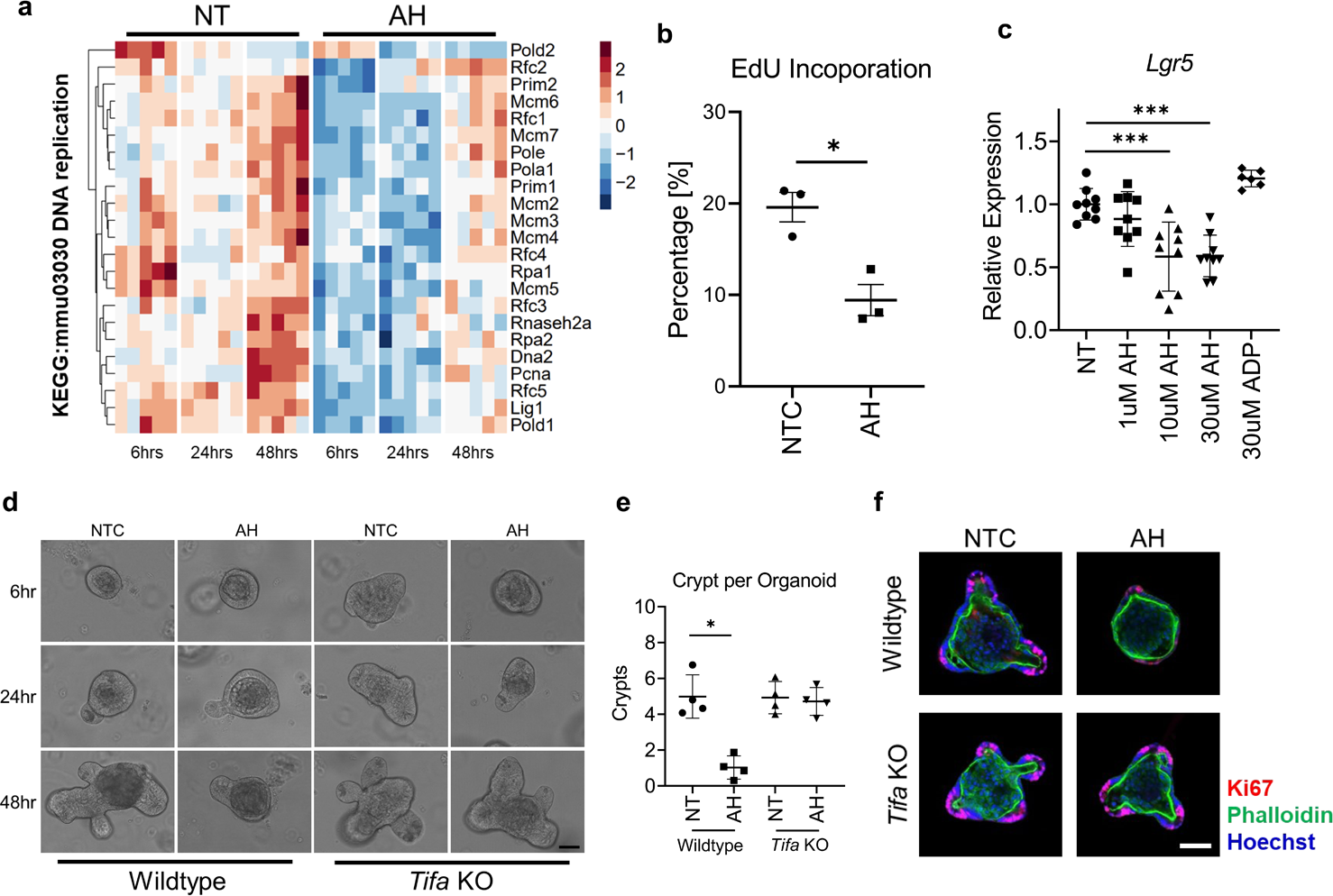
ADP-Hep induces a TIFA-dependent blunting of IEC proliferation. **a,** Heatmap generated from bulk RNA-seq data of wildtype ileal organoids untreated or treated with ADP-Hep for 6, 24 or 48 hours. Transcripts are from the KEGG term: DNA replication. **b,** Percent cells positive for EdU incorporation after 48 hours of ADP-Hep stimulation and 2 hours of EdU incorporation, as assessed by flow cytometry (n = 3). **c,** RT-qPCR of ileal organoids stimulated with ADP-heptose at 1-30µM and ADP at 30µM (n ≥ 6). **d,** Representative images from bright field microscopy of organoids treated with ADP-Hep [20µg/mL] for 6, 24 or 48 hours (n = 4). **e,** Quantification of number of crypts per organoid of organoids stimulated with ADP-Hep [20µg/mL] for 48 hours (n = 4, >10 organoids quantified per biological rep). **f,** Representative image of immunofluorescent staining for Ki67 (red) in wildtype or *Tifa^-/-^* organoids treated with ADP-Hep [20µg/mL] for 48 hours (n = 3). ***P<0.001, *P<0.05. Scale bar 50µm (d), 50µm (f).

## ADP-Hep inhibits proliferation through apoptosis

Since *Alpk1* is expressed in ISCs, we next hypothesized that ADP-Hep may blunt epithelial proliferation and organoid growth by killing ISCs. Immunoblot analysis of murine organoids treated with ADP-Hep showed the accumulation of the active fragments of caspase-8 (p18) and caspase-3 (p12/p17), and the cleavage of Parp-1 (target of active caspase-3), in a *Tifa*-dependent manner (Fig. 3a). Additionally, ADP-Hep-treated organoids did not show accumulation of the pyroptosis-associated p30 fragment of Gsdmd (Supplementary Fig. 2a) or the necroptosis-associated phospho-Mlkl (Supplementary Fig. 2b), indicating that ADP-Hep specifically induced apoptosis and not pyroptosis nor necroptosis. Similarly, human intestinal organoids stimulated with ADP-Hep showed induction of NF-κB signalling and caspase-dependent apoptosis (Fig. 3b). Immunofluorescent staining for cleaved caspase-3 of WT murine organoids treated with ADP-Hep for 4 hours revealed the accumulation of apoptotic cells within the crypts, which was not observed in untreated organoids (Fig. 3c, white arrows). These organoids also exhibited strong caspase-3 positive signal inside the crypt lumen (Fig. 3c, white arrow heads), likely resulting from the shedding and accumulation of apoptotic cells and apoptotic debris into this compartment.

**Figure 3.**
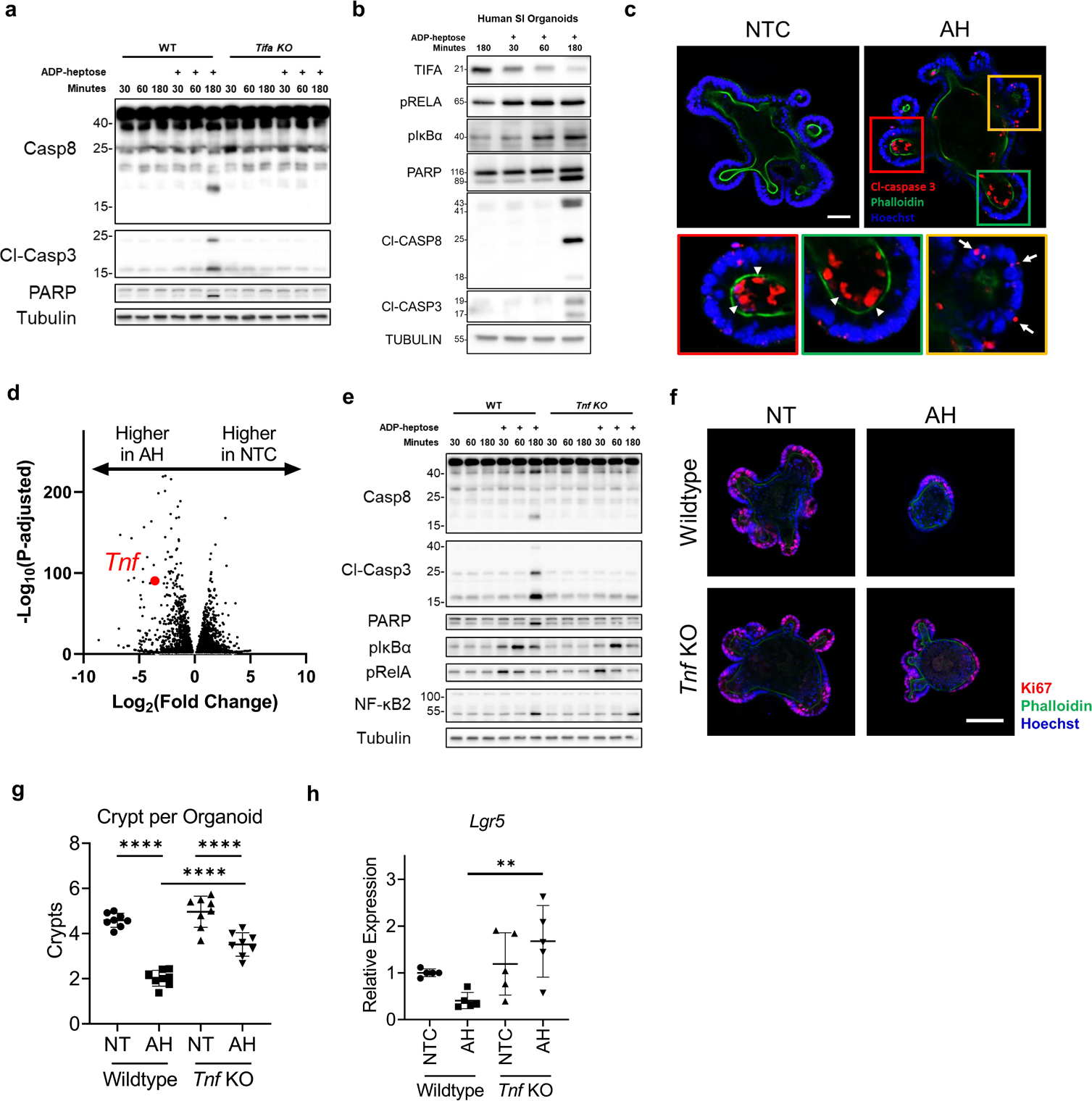
ADP-Hep induces a TIFA- and TNF-dependent blunting of IEC proliferation through apoptosis. **a,** Representative immunoblot of ileal organoids from either wildtype or *Tifa^-/-^* mice, stimulated with ADP-Hep for 30, 60 or 180 minutes (n = 3). **b,** Representative immunoblot of human ileal organoids stimulated with ADP-Hep for 30, 60 or 180 minutes (n = 3). **c,** Representative immunofluorescent staining for cleaved caspase 3 (red) in wildtype organoids treated with ADP-Hep [20µg/mL] for 4 hours (n = 3). **d,** Volcano plot of differential mRNA expression of ileal organoids stimulated with ADP-heptose [20µg/mL] for 6 hours compared to untreated. **e,** Representative immunoblot of ileal organoids from either wildtype or *Tnf^-/-^* mice, stimulated with ADP-Hep for 30, 60 or 180 minutes (n=3). **f,** Representative immunofluorescence staining of wildtype or *Tnf*^-/-^organoids treated with ADP-Hep [20µg/mL] for 48 hours (n = 3). **g,** Quantification of number of crypts per organoid of organoids stimulated with ADP-Hep [20µg/mL] for 48 hours (n = 8, >10 organoids quantified per biological rep). **h,** RT-qPCR of ileal organoid generated from either wildtype or *Tnf^-/-^* mice, treated with ADP-Hep for 48 hours (n = 5). ****P<0.0001, **P<0.01. Scale bar 40µm (c), 100µm (f).

The ADP-Hep-dependent cleavage of caspase-8 suggested the likely involvement of extrinsic apoptosis. Interestingly, our bulk RNAseq data revealed that *Tnf* was strongly upregulated (12-fold) by ADP-Hep treatment (Fig. 3d), which prompted us to hypothesize that ADP-Hep-induced apoptosis may be caused by Tnf-dependent paracrine signalling. In agreement, ADP-Hep-dependent accumulation of cleaved caspase-3, cleaved caspase-8 and cleaved Parp-1 was fully abrogated in organoids from *Tnf^-/-^* mice, while NF-κB induction was normal (Fig. 3e). Since Tnf is a known NF-κB transcriptional target, these data together suggest that Tnf-dependent apoptosis triggered by ADP-Hep likely occurred in a second wave of signalling following NF-κB induction and required *Tnf* upregulation. In support of this, ADP-Hep-dependent apoptosis was blunted in WT organoids treated with an anti-Tnf neutralizing antibody (Supplementary Fig. 2c). Interestingly, while Tnf was necessary for ADP-Hep-dependent apoptosis, it was insufficient on its own to drive apoptosis in WT organoids (Supplementary Fig. 2c), implying that ADP-Hep not only upregulates Tnf but also sensitizes crypt IECs to Tnf-mediated apoptosis.

To assess if Tnf-driven apoptosis was causal in mediating stunting of epithelial proliferation and crypt architecture in ADP-Hep-treated organoids, we next performed immunofluorescent staining on organoids derived from WT and *Tnf^-/-^* littermate mice. Ki-67 staining of ADP-Hep-treated organoids showed significant preservation of crypt structure and numbers in *Tnf^-/-^* organoids as compared to WT organoids (Fig. 3f and quantifications in Fig. 3g). Moreover, while ADP-Hep blunted *Lgr5* expression in WT organoids, this effect was not observed in *Tnf^-/-^* organoids (Fig. 3h). The preservation of crypt numbers in *Tnf^-/-^* organoids, however, was partial since their numbers remained significantly lower in *Tnf*^-/-^ organoids stimulated with ADP-Hep as compared to untreated organoids (Fig. 3g). This suggests the existence of additional mechanisms, besides Tnf-dependent apoptosis induction, by which ADP-Hep inhibits proliferation of the intestinal epithelium. Together, these data demonstrate that ADP-Hep disrupts crypt architecture and inhibits proliferation of the epithelium, in part through induction of apoptosis within the crypts, in a Tnf-dependent manner.

## ADP-Hep induces revSCs to reconstitute the ISC pool

While our results so far have shown that ADP-Hep treatment induces acute loss of cells in the crypt, blunted IEC proliferation, loss of crypt architecture and reduction of homeostatic ISC markers, we speculated that, upon removal of ADP-Hep, the intestinal epithelium may recover through a regenerative program. Indeed, in organoids treated for 24 hours with ADP-Hep followed by a media change without ADP-Hep and allowed to grow for another 24 hours, we observed the budding of new crypts at the bottom of which active Ki67^+^ cells were found (Fig. 4a), suggesting that a regenerative stem cell program was initiated. Interestingly, our bulk RNAseq dataset of ADP-Hep-treated organoids revealed that *Ly6a* and *Clu*, two of the main markers of revSCs, were strongly upregulated (Fig. 4b-d). *Ly6a* induction was more pronounced at the 6-hour time point, while *Clu* upregulation appeared to be more progressive, being the highest at 48 hours; at this time point, *Clu* was the third most significantly differentially-expressed gene in our dataset (Fig. 4d). In addition, genes of the *Anxa* family, which are also signatures of revSCs ^7,8^, were massively upregulated by ADP-Hep treatment (Supplementary Fig. 2a-c). More broadly, the revSC induction was previously shown to be dependent on the YAP/TAZ pathway and on YAP-dependent suppression of Wnt signalling ^5^. In agreement, ADP-Hep treatment upregulated YAP/TAZ-dependent genes as early as 6 hours post treatment (Fig. 4e), while the majority of Wnt target genes was concomitantly downregulated (Fig 4f). Importantly, revSC induction required ADP-Hep-dependent triggering of apoptosis, since *Clu* induction was fully abrogated in organoids derived from *Tnf^-/-^* mice (Fig. 4g).

**Figure 4.**
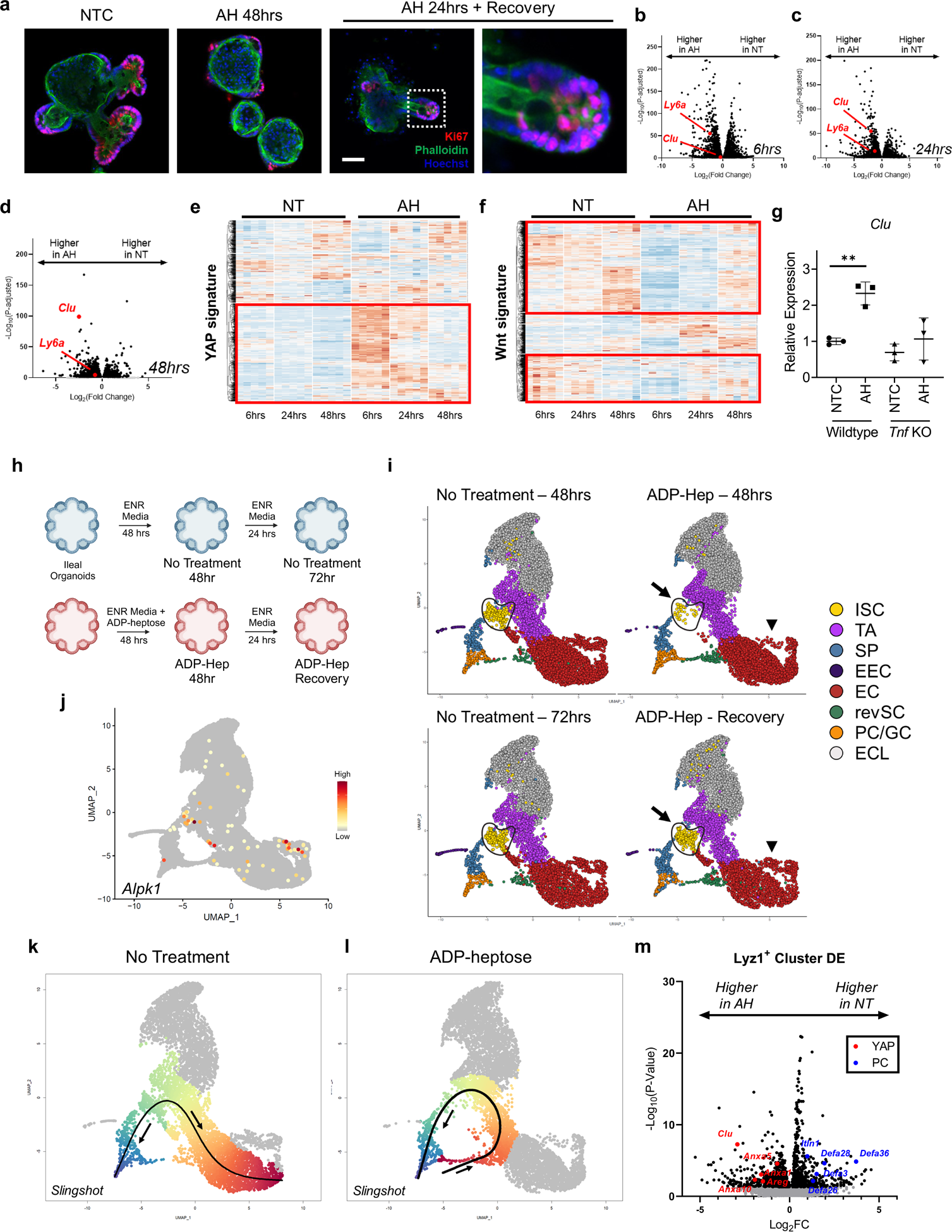
ADP-Hep induces a regenerative program to reconstitute ISCs. **a,** Representative immunofluorescent staining of wildtype organoids untreated or treated with ADP-Hep [20µg/mL] for 48 hours or treated for 24 hours with ADP-Hep followed by recovery in ENR media for 24 hours (n = 3). **b-d,** Volcano plot of differential mRNA expression of ileal organoids stimulated with ADP-Hep [20µg/mL] for 6, 24 and 48 hours. **e-f,** Heatmap generated from bulk RNA-seq data of wildtype ileal organoids treated with ADP-Hep for 6, 24 or 48 hours. Signatures are from Gregorieff A et al. Nature. 2015. **g,** RT-qPCR of ileal organoid generated from either wildtype or *Tnf^-/-^* mice, treated with ADP-Hep for 48 hours. **h,** Schematic of single cell experiment conducted on murine ileal organoids. **i,** UMAP plot of untreated, ADP-Hep treated, and ADP-Hep treated and recovered organoids. **j,** Expression of *Alpk1* overlayed on UMAP plot. **k-l,** Trajectory analysis of untreated and ADP-Hep treated organoids. **m,** Volcano plot generated from differentially expressed genes between untreated and ADP-Hep treated, Lyz1+ cells. Cells subset from scRNA-seq organoid dataset. **P<0.01.

To provide a deeper characterization of the acute insult induced by ADP-Hep and of the subsequent epithelial regeneration in the recovery phase, we performed single-cell RNAseq (scRNAseq) analysis of WT intestinal organoids either stimulated for 48 hours with ADP-Hep or 48 hours + 24 hours of recovery, as well as their respective pairwise matching untreated controls at 48 and 72 hours (Fig. 4h). Using the R package Seurat, we integrated and clustered a total of 27,207 cells at an average read depth of 47k/cell and visualised these populations using uniform manifold approximation and projection (UMAP; Fig. 4i). The analysis allowed defining specific cell population clusters for: ISCs, transit amplifying cells, secretory progenitors, differentiated Paneth and goblet cells, enteroendocrine cells and enterocytes, as highlighted by the expression profile of common cell population markers ^13–16^ (Supplementary Fig. 4). Of note, the clustering was not able to differentiate mature Paneth and goblet cells into distinct clusters, in line with results from a previously published scRNAseq organoid dataset ^16^. Additionally, our dataset revealed the presence of previously reported enterocyst-like cells (ELCs) ^14–16^ (Fig. 4i, grey cells), which are found as spherical organoids that only differentiate into enterocytes (Supplementary Fig. 5a) as further supported by Slingshot trajectory analysis of these ELCs predicting differentiation exclusively into enterocytes (Supplementary Fig. 5b). Finally, our dataset also showed the presence of a small constitutive cluster of revSCs, marked by *Clu, Ly6a* and *Anxa1* (Fig. 4i; Supplementary Fig. 4), which confirms previously published work on intestinal organoids ^14–16^. It is likely that this constitutive revSC cluster represents a particularity of organoid culturing since we did not identify constitutive revSCs in our previous scRNAseq analysis of crypt-enriched IECs from murine ileal tissue ^13^.

The most remarkable effect that we observed through our scRNAseq analysis was the severe blunting of the ISC pool, marked by *Lgr5*, *Olfm4*, *Lrig1* and *Mki67*, at 48 hours of ADP-Hep treatment and its full reconstitution in the 48 hours + 24 hours of recovery condition (Fig. 4i, arrow; Supplementary Fig. 4). Importantly, *Alpk1* expression was restricted to the ISC pool that was ablated by ADP-Hep, and additionally to a minor subset of differentiated enterocytes that were also lost during ADP-Hep treatment (Fig. 4i, arrowhead and Fig. 4j). The expression of *Alpk1* in organoid ISCs further confirms our previous scRNAseq ^13^ and RNA-ISH data obtained on ileal tissue (see Fig. 1). This acute loss of ISCs is also in line with our data (see Figs. 2 and 3) showing blunted proliferation and crypt architecture disruption in ADP-Hep-treated organoids, and further suggests that these effects were a consequence of the specific ablation of the ISC pool.

The acute and transient loss of ISCs in response to ADP-Hep, revealed by our scRNAseq analysis, offers a unique opportunity to dissect the mechanisms by which ISCs are rederived following acute injury. In particular, a key unanswered question is the identity of the cells from which the revSCs derive following injury. Answering this fundamental question is challenging with the current models of revSC induction (such as γ-irradiation or dextran sodium sulfate (DSS)-mediated epithelial injury ^6^) because the insult likely damages the epithelium with little selectivity at multiple places simultaneously. In contrast, ADP-Hep-mediated ISC ablation appears to be highly selective, which facilitates the analysis of the mechanisms underlying the emergence of revSCs. Through Slingshot trajectory analysis, we first observed that under control conditions, constitutive ISCs gave rise to secretory cells and enterocytes, as expected (Fig. 4k). However, cell differentiation trajectories appeared strikingly altered in ADP-Hep-treated conditions with the predicted origin of the trajectory being located at the boundary of the *Lyz1*^+^ Paneth/Goblet cluster and the constitutive revSC cell cluster (Fig. 4l). Further differential gene expression analysis of the extracted *Lyz1*^+^ cluster between untreated and ADP-Hep-treated organoids showed induction of transcripts associated with YAP signalling, including *Clu, Anxa1, Anxa5, Anxa10* and *Areg*. Moreover, gene ontology (GO) term enrichment analysis of the *Lyz1*^+^ cluster treated with ADP-Hep revealed enrichment for the GO term Mitotic cell cycle (GO: 0000278). Concurrently, transcripts associated with differentiated Paneth cells, including *Itln1, Defa28, Defa36, Defa2* and *Defa26*, were reduced in the ADP-Hep-treated *Lyz1*^+^ cluster (Fig. 4m), while those associated with mature goblet cells (*Muc2* and *Tff3*) were unchanged. This suggests that the emergence of *Clu*^+^ cells in the *Lyz1*^+^ cluster is associated with a specific de-differentiation of Paneth cells rather than goblet cells. In support, sub-clustering of the Paneth/Goblet population revealed that *Clu*, *Ly6a* and *Anxa1* transcripts, which are induced by ADP-Hep, were found predominantly within a Paneth cells subcluster (subcluster 3; Supplementary Fig. 7).

Together, these data demonstrate that ADP-Hep induces a revSC program in IECs that reestablishes the ISC pool lost by apoptosis, and suggest that the *Lyz1*^+^ Paneth cell population likely contains the cells from which revSCs emerge upon ADP-Hep-mediated injury.

## *Lyz1*^+^ Paneth cells serve as cells of origin for revSCs following ADP-heptose treatment

Next, we aimed to provide direct evidence of the role played by Paneth cell de-differentiation in the emergence of revSCs following ADP-Hep treatment, and to visualize cells in their transitionary state between Paneth cells and revSCs, the existence of which being predicted by our scRNAseq data. Interestingly, YAP/TAZ signalling is critical for epithelial regeneration following injury and our scRNAseq data revealed the expression of YAP target genes in Paneth cells following ADP-Hep treatment (see figure 4l). This was a surprising finding given the fact that YAP expression is normally low in Paneth cells and inhibits the differentiation towards the Paneth cell lineage ^5,17^. Thus, to confirm our scRNAseq data at the protein level, ADP-Hep-treated organoids were first imaged by immunofluorescence using an anti-YAP antibody. This revealed a significant upregulation of nuclear YAP, which is indicative of YAP activation, in a subset of Paneth cells following ADP-Hep treatment (Fig. 5a, b). This nuclear localization of YAP was specific to Paneth cells as the ISCs intercalated between Paneth cells did not show this effect (Fig. 5c). Next, using intestinal organoids derived from *Clu* EGFP reporter (*Tg(Clu-EGFP)OD95Gsat*) mice, we performed immunofluorescent staining following ADP-Hep stimulation, which revealed cells co-positive for the Paneth cell marker Itln1 and for EGFP (Fig. 5d), an event not observed in untreated conditions. Therefore, these imaging approaches allowed for the identification of cells likely transitioning from Paneth cells to revSCs following ADP-Hep treatment.

**Figure 5.**
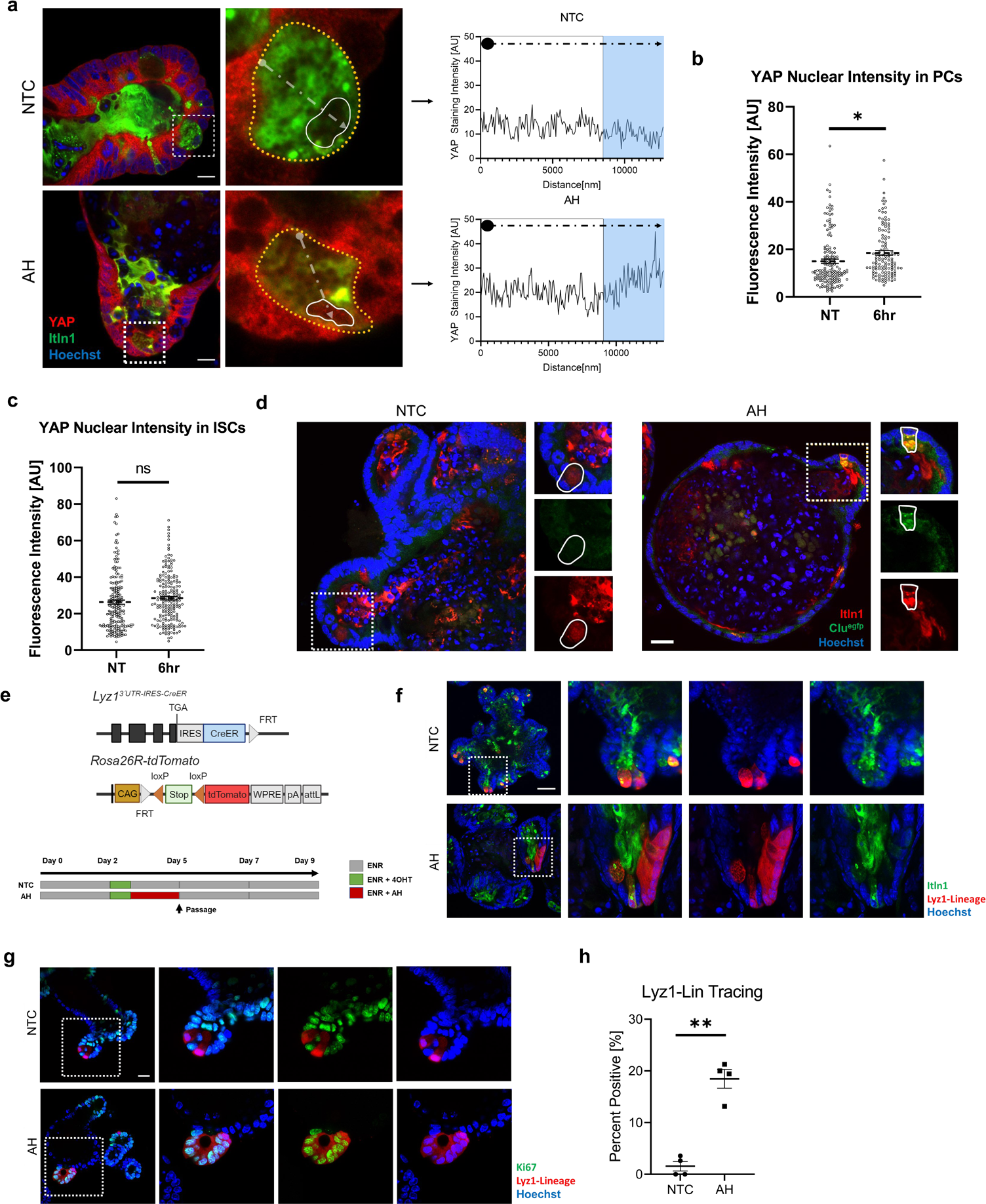
ADP-Hep induces a regenerative program to rederive ISCs from PC dedifferentiation. **a,** Left, representative image of immunofluorescent staining of YAP (red) and Itln1 (green) of unstimulated and ADP-Hep stimulated organoids at 6 hours. Right, quantification of fluorescent intensity. Arrows indicate path of quantification. **b,** Violin plot of quantification of YAP nuclear intensity from Itln1^+^ cells (quantification of 50 cells from 4 biological replicates each). **c,** Violin plot of quantification of YAP nuclear intensity of ISC intercalating Itln1^+^ cells (quantification of 50 cells from 4 biological replicates each). **d,** Representative image of immunofluorescent staining of Itln1(red) and Clu-EGFP (green) on organoids from *Clu* reporter mice stimulated with ADP-Hep. **e,** Schematic of Lyz1 lineage tracing mice and experimental outline for PC lineage tracing experiment. **f,** Representative image fluorescent staining for Itln1 (green) from organoids generated from Lyz1 lineage tracking mice treated with ADP-Hep (n = 3). **g,** Representative image of fluorescent imaging of Ki67 (green) from organoids generated from Lyz1 lineage tracking mice treated with ADP-Hep (n = 3). **h,** Quantification of percent ribbon positive organoids from either unstimulated or ADP-Hep stimulated organoids (n = 4). **P<0.01, *P<0.05. Scale bar 10µm (a), 20µm (d), 50µm (f), 20µm (g).

If Paneth cell de-differentiation to revSC following ADP-Hep treatment is functional, we would expect to observe clonal expansion of cells originating from the Paneth cell lineage that progressively migrate upwards from the crypt base. To assess this, we generated intestinal organoids from *Lyz1*^+^ lineage tracing reporter (*Lyz13′UTR-IRES-CreER; Rosa26R-tdTomato*) mice (Fig. 5e), which were recently generated and characterized ^18^. In untreated organoids, the tdTomato marker was almost exclusively co-localized with Itln1 (Fig. 5f, top and quantification in Fig. 5h), thus confirming that the lineage tracing was not “leaky” and remained restricted to Paneth cells. In contrast, ADP-Hep treatment induced the formation of tdTomato^+^ ribbons, the majority of which showed Itln1-negative staining (Fig. 5f bottom, quantification in Fig. 5h), thereby demonstrating that cells originating from the Lyz1 lineage acquired pluripotency in response to ADP-Hep. In further support, while cells of the Lyz1 lineage were negative for Ki67 in the untreated condition, which is expected as Paneth cells are post-mitotic, Lyz1 lineage-traced cells displayed robust Ki67 staining following ADP-Hep treatment, thus showing that these cells had acquired the ability to proliferate (Fig. 5g).

Therefore, ADP-Hep induces revSC and epithelial regeneration through Paneth cell de-differentiation.

## Induction of revSCs *in vivo* is dependent on Tifa signalling

Since ADP-Hep is an intermediate of LPS biosynthesis and is thus ubiquitous among gram-negative bacteria, it is likely that this molecule would be abundantly found in the intestinal lumen, being produced by the resident microbiota. To confirm this, we performed high-performance liquid chromatography/mass-spectrometry analysis of stool samples collected from specific pathogen-free (SPF) versus GF mice, which revealed that ADP-Hep was present in samples from SPF but not GF mice (Fig. 6a). While ADP-Hep is constitutively present in the gut, we reasoned that, in resting conditions, this bacterial metabolite would have limited access to the bottom of intestinal crypts as bacteria are normally excluded from this niche, thereby protecting ISCs from ADP-Hep-mediated apoptosis and induction of revSC programs. We further hypothesized that upon epithelial injury, such as in the case of γ-irradiation, disruption of the intestinal crypt architecture would allow ADP-Hep to gain access to the stem cell zone to initiate a revSC program.

**Figure 6.**
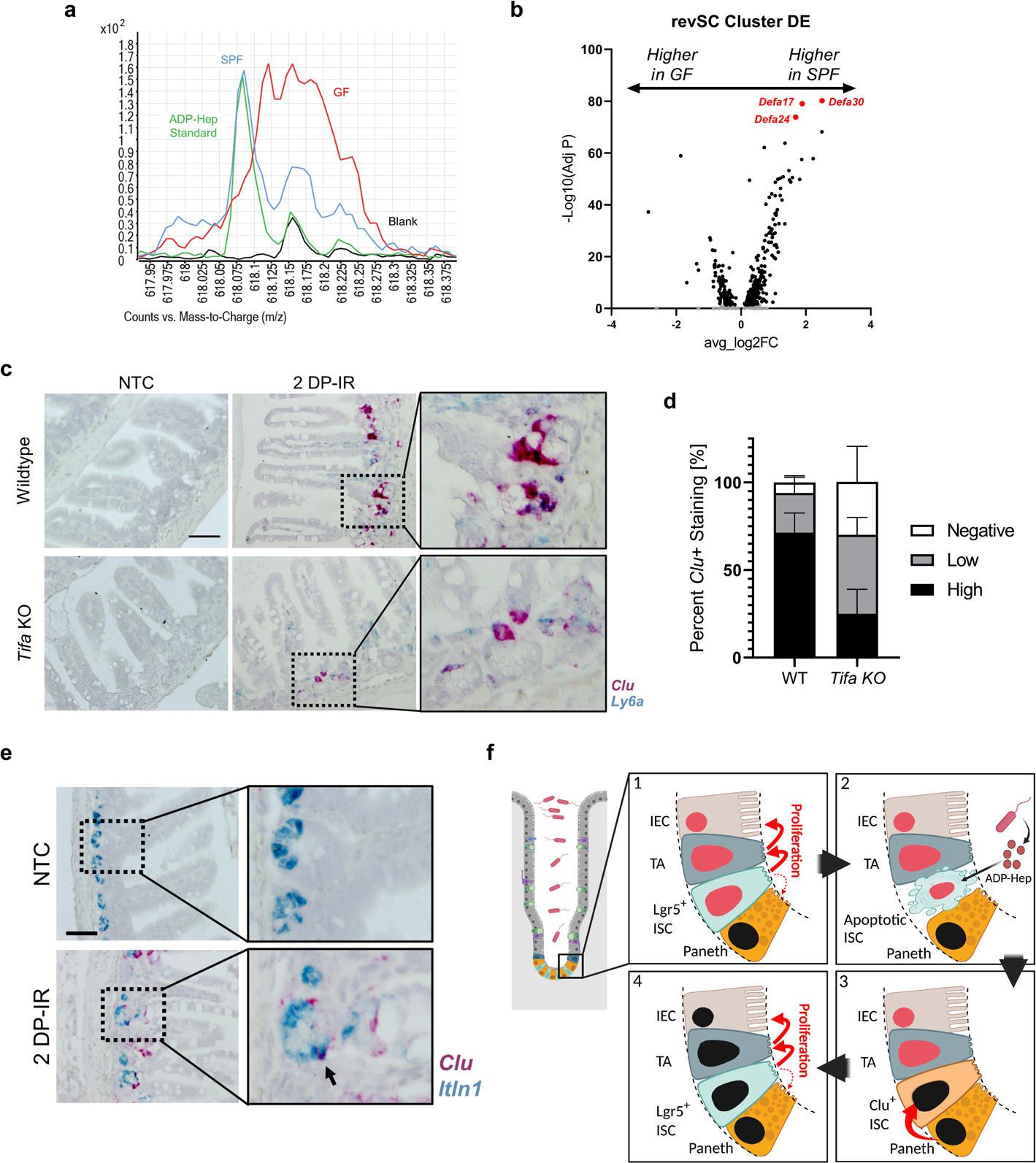
Induction of revSCs *in vivo* is dependent on Tifa signalling. **a,** Retention time from mass spectrum analysis of stool pellets pooled from either GF or SPF mice and an ADP-Hep standard (n = 3). **b,** Volcano plot of differential expression analysis from scRNA-seq dataset of crypt enriched IECs, comparing revSCs transcription between irradiated SPF and GF mice. **c,** Representative dual RNA-ISH for *Clu* (red) and *Ly6a* (blue) from either WT or *Tifa^-/-^* mice two days post irradiation (n ≥ 4). **d,** Quantification of frequency of crypts from panel c for either negative, low or high for *Clu* staining in panel c (n ≥ 4, 100 crypts quantified per mouse). **e,** Dual RNA-ISH for *Clu* (red) and *Itln1* (blue) from WT mice two days post irradiation (n = 4). **f,** Graphical model of ADP-Hep detection by ISCs and the subsequent reversion of PCs to ISCs. Scale bar 50µm (c), 50µm (e).

To test this hypothesis, we first γ-irradiated SPF and GF mice, collected crypt-enriched epithelial ileal tissue post-irradiation and performed scRNAseq. While the full analysis of this dataset will be described elsewhere (Tsang et al., manuscript in preparation), we focussed here on the *Clu*^+^ population. Interestingly, differential expression analysis of *Clu*^+^ revSCs showed significantly higher expression of Paneth cell-associated transcripts, including *Defa30* and *Defa24*, in revSCs from SPF as compared to GF mice (Fig. 6b). This suggests that revSCs may have arisen from Paneth cells in this model of irradiation-induced epithelial damage, and that this event may be enhanced by microbiota-associated factors. To further assess if ADP-Hep and the Alpk1-Tifa pathway were involved in this process, WT and *Tifa*^-/-^ littermate mice were γ-irradiated and ileal tissue was collected 2 days post-irradiation (2dpi) for RNA-ISH. While *Clu* and *Ly6a* expression were strongly upregulated by irradiation in WT mice, as previously reported ^6^, this induction was significantly blunted in *Tifa*^-/-^ mice as compared to WT mice (Fig. 6c, d). Moreover, RNA-ISH demonstrated frequent colocalization of *Lyz1* and *Clu* (Fig. 6e), thus providing further support to our organoid data by suggesting that Paneth cell de-differentiation may contribute to revSC induction in this model of irradiation. Together, these data demonstrate that Tifa-dependent signalling plays a critical role in amplifying revSC induction *in vivo* following γ-irradiation.

The role played by the microbiota in the context of intestinal inflammation and barrier disruption is definitely complex. While it is established that bacterial translocation to the intestinal mucosa can fuel inflammation and amplify barrier destruction in a vicious circle, microbe-dependent protective mechanisms also exist, which rely either on specific metabolites (e.g., short chain fatty acids, indole derivatives) or innate immune pathways of microbial detection (e.g., NOD2 signaling whose dysregulation is associated with CD) ^1,2^. The Alpk1-Tifa pathway of revSC induction following epithelial injury provides a novel perspective on the potential impact of microbial metabolites on ISC function and intestinal barrier protection. Indeed, in contrast to the idea that the ISC pool needs to be protected at all costs to avoid disruption of the intestinal epithelial architecture and barrier breach, our results suggest that innate immunity may have selected a defense mechanism through which the detection of ADP-Hep, a common MAMP produced by Gram-negative bacteria, initiates a rapid “reset” of the ISC pool through apoptosis induction and revSC-dependent restoration of homeostasis. It remains to be determined if the ADP-Hep-dependent process of revSC induction is a pathogenic or a protective mechanism. While we favor the latter model because ADP-Hep is a common MAMP detected by the host innate immune Alpk1-Tifa pathway, it remains unclear why the acute ablation of homeostatic ISCs followed by the rapid emergence of revSCs would be protective in the context of infection with ADP-Hep-expressing bacteria. An interesting hypothesis could be that this process has been selected to limit the risk that genotoxic bacteria reaching the bottom of intestinal crypts would induce DNA damage in the ISC pool, thereby provoking fixation of potentially deleterious mutations in the intestinal epithelium. Indeed, genotoxic microorganisms (including colibactin-expressing *Escherichia coli* ^19^ and *Fusobacterium nucleatum* ^20^) are frequently found within the class of Gram-negative bacteria that produce ADP-Hep. Interestingly, mutations in *ALPK1* were associated with Lynch-like syndrome, in which patients meet the clinical criteria for Lynch syndrome (the most common cause of hereditary colorectal cancer (CRC)) but do not present with typical mutations in DNA mismatch repair genes ^21^. Moreover, *ALPK1* expression was found to be reduced in CRC tumours as compared to normal adjacent tissue, suggesting that it acts as a tumour suppressor in CRC ^22^. Further studies are thus required to delineate the role of ADP-Hep detection by the Alpk1-Tifa pathway in ISCs, and the impact of the induction of a revSC program, in the protection against enteric bacterial infection and possibly against CRC.

## Material and Methods

### Reagents for treatments

Reagents used for treatment are as listed: ADP-heptose (tlrl-adph-l, InvivoGen), TNF (315-01A-5UG, Cedarlane), Z-VAD-FMK (ALX-260-020-M0001, Enzo Life Sciences), SM164 (B1816-250, BioVision), Lipoteichoic acid (tlrl-slta, InvivoGen), Lipopolysaccharide (14011S, CST), Pam3CSK4 (tlrl-pms, InvivoGen), L18-MDP (tlrl-lmdp, InvivoGen) C12-iE-DAP (tlrl-c12dap, InvivoGen), and Flagellin (tlrl-stfla, InvivoGen).

### Murine Organoid Culture

Organoids were generated from murine intestine as described ^23^. In brief, crypts per isolated from murine intestine and cultured in domes of Cultrex Pathclear Reduced Growth Factor Basement Membrane Extract (3533-005, R&D Systems), covered by ENR media comprised of Advanced DMEM/F12 (12634028, Gibco), 1x GlutaMAX (35050061, Gibco), 1x Penicillin-Streptomycin (450-201-EL, Wisent), 10mM HEPES (15630080, Gibco), 1x B27 (17504044, Gibco), 1x N2 (17502048, Gibco), 50ng/mL mEGF (PMG8043, Gibco), 1mM N-acetylcysteine (A9165-100G, Sigma), 10% v/v R-spondin conditioned media (in-house) and 10% v/v Noggin conditioned media (in-house). Organoids were grown in ENR with 10μM Y-27632 for 2 days post passage, and without Y-27632 there after. Organoids were maintained in a 5% CO_2_ incubator at 37°C. For comparisons between wildtypes and knockouts, organoids were generated from litter mate mice.

### Human Organoid Culture

Human organoids were prepared from the small intestine of a human organ donor diverted to research by the Trillium Gift of Life Network in accordance with human bioethics protocol to H.Y.G. Human organoids were generated as described ^24^. In brief, fresh small intestinal tissue was digested in 2 mg/mL Collagenase Type I (17100017, Thermo Fisher) and broken down into smaller fragments by mincing with scissors followed by pipetting. The tissue was digested for 40 minutes at 37°C. After digestion, crypts were dissociated from the tissue with vigorous pipetting. Crypts were filtered out of the tissue sample with a 70uM cell strainer and pelleted at 300 x g for 5 minutes at 4°C. The pellet was resuspended in Cultrex Pathclear Reduced Growth Factor Basement Membrane Extract (3533-005, R&D Systems) and covered by human small intestinal plating media comprised of 50% v/v L-WRN Conditioned Media ^25^, 5% v/v R-Spondin 1 Conditioned Media (in-house), 35% v/v Base Media [Advanced DMEM/F12 (12634028, Gibco), 1x Penicillin-Streptomycin (450-201-EL, Wisent), 20% Fetal Bovine Serum (080-150, Wisent)], 1x GlutaMAX (35050061, Gibco), 10mM HEPES (15630080, Gibco), 1x B27 (17504044, Gibco), 1x N2 (17502048, Gibco), 10mM Nicotinamide (72340-100G, Sigma), 500nM A-8301 (SML0788-5MG, Sigma), 10uM SB202190 (S7067-5MG, Sigma), 50ng/mL Human Recombinant EGF (AF-100-15, Preprotech), 1mM N-acetylcysteine (A9165-100G, Sigma), 10nM [Leu15]-Gastrin I Human (G9145, Sigma), 10uM Y-27632 (1254, Tocris), and 250nM CHIR 99021 (4423, Tocris). Organoids were grown in a 37°C/5% CO2 incubator for 7 days. Plating media was replenished every 3 days. For human organoid maintenance, plating media was used for 3 days post passage, and then grown without Y-27632 and CHIR 990221 thereafter.

### RNA isolation and sequencing

RNA was isolated from ADP-heptose stimulated organoids using the GeneJET RNA purification kit (K0732, Thermo Scientific) according to the manufacturer’s instructions. RNA samples were prepared, sequenced, and analyzed by Novagene. In brief, samples were sequenced at 20 million reads per sample 150bp paired end using an Illumina Novaseq 6000. Alignment was performed using STAR software (v 2.6.1d), differential analysis was performed using DESeq2 (v 2.6.1d) and ClusterProfiler (v 3.8.1) was used for enrichment analysis. Heatmaps were generated using pHeatmap (v 1.0.12).

Previously published RNA-sequence datasets were accessed from NCBI GEO: GSE81125 and NCBI GEO: GSE60163. Analysis of GSE81125 was performed using edgeR (v 3.34.1).

### RNA in situ hybridization

RNA *in situ* hybridization was performed using the RNAScope 2.5 HD Duplex Assay kit (ACD) according to the manufacturer’s instructions. In brief, tissue was isolated and fixed in 10% neutral buffered formalin (HT501128-4L, Sigma-Aldrich) for 48 hours before being embedded in paraffin. Paraffin embedded tissue was sectioned at a thickness of 5μm.

### Mouse lines

*Tifa* knockout mice were generated using CRISPR/Cas9 by the Model Production Core at The Centre for Phenogenomics (TCP). All procedures involving animals were performed in compliance with the Animals for Research Act of Ontario and the Guidelines of the Canadian Council on Animal Care (CCAC). The TCP Animal Care Committee reviewed and approved all procedures conducted on animals at TCP. Three synthetic gRNAs were co-injected into C57Bl6/NCrl zygotes to target the single coding exon of mouse Tifa common to all transcripts. Two target the 5’ end of the coding region (gRNA-U5 and gRNA-U3) and one targets the 3’ end of the coding region (Supplementary Table 1). These gRNA sequences were scored for specificity using http://crispr.mit.edu/ and were predicted to have no off targets with less than 3 mismatches. *Tnf* knockout mice were obtained from the Jackson Laboratory (Strain #007914). *Clu* EGFP reporter (*Tg(Clu-EGFP)OD95Gsat*) mice were obtained from a previous study ^6^. Paneth cell lineage tracing (Lyz1^3′UTR-CreER^;R26-tdTomato) mice were obtained from a previous study ^18^. All mice were housed under SPF conditions at the Division of Comparative Medicine, University of Toronto. Adult mice (8-20 weeks) littermate wildtype and knockout mice were used for organoid generation and *in* vivo experimentation. All mice experiments were approved by the local Animal Ethics Review Committee. Genotyping primers for mice used can be found in Supplementary Table 2.

### RT-qPCR

Isolated RNA (described above) was treated with DNAase I (EN0521, Thermo Scientific) according to manufacturer’s instructions. Reverse transcription was performed using the high-capacity cDNA reverse transcription kit (4368814, Applied Biosystems) according to manufacturer’s instructions. Primers used are shown in Supplementary Table 3.

### Immunoblotting

Organoids were lysed in RIPA buffer and centrifuged at 10,000*g* for 10 minutes followed by boiling with Laemmli blue loading buffer. Samples were run on acrylamide gels and transferred onto PVDF membranes. Membranes were blocked 5% milk or 2.5% BSA (for phosphor-proteins) in TBS-T and incubated with the following antibodies: Phospho-IκBα (2859S, CST, 1/1000), Phospho-NF-κB p65 (ab32518, Abcam, 1/1000), NF-κB2 p100/p52 (11948S, CST, 1/1000), α/β-Tubulin (2148S, CST, 1/1000), Cleaved Caspase-8 (8592S, CST, 1/1000), Cleaved Caspase-3 (9661S, CST, 1/1000), PARP (9542S, CST, 1/1000), TIFA (61358, CST, 1/1000), GSDMD (ab209845, Abcam, 1/1000), MLKL (37705S, CST, 1/1000) and phospho-MLKL (37333S, CST, 1/1000).

### Immunofluorescence staining

Immunofluorescence on organoids was performed as previously described ^23^. Antibodies used are as follows: Cleaved Caspase-3 (9661S, CST, 1/100), Ki67 (12202S, CST, 1/200), GFP (A-11122, Invitrogen, 1/200), Intelectin-1 (MAB8074, Bio-Techne, 1/100), YAP (14074S, CST, 1:100).

### Single-cell RNA sequence analysis of organoids

Stimulated organoids were dissociated as described in Bues et al, Nature Methods ^14^. In brief, single-cell dissociation buffer was prepared in PBS (311-425-CL, Wisent) with, 10 mg per ml Bacillus licheniformis protease (P5380-100MG, MilliporeSigma), 5 mM EDTA (EDT111.100, Bioshop), 5 mM EGTA (2807, Toris), 10 µg per ml DNase I (11284932001, MilliporeSigma) and 0.68X Accutase (A6964, MilliporeSigma). Organoids were incubated in single-cell dissociation buffer for 15 minutes at 37°c with periodic disruption by pipetting. An estimated 10,000 cells were collected for BD Rhapsody single-cell isolation and library preparation following the manufacture’s instructions. Sequencing was performed on an Illumina Novaseq 6000. Alignment to mm10/GRCm38 was performed using the BD Rhapsody Whole Transcriptome Analysis (WTA) pipeline ^26^.

In R 4.1.0, Seurat ^27^ (4.1.0) was used for analysis of single-cell expression matrices. Cells with less than 5000 reads, 1000 features or >50% mitochondrial genes were filtered out. Feature were log normalised and the top 2000 variable genes were selected for dataset integration. We additionally used Seurat to perform PCA dimensionality reduction and identify cell clusters. Cluster cell type labels were assigned based off previously published cell markers in organoid and the intestinal epithelium ^13–16^. Uniform manifold approximation and projection (UMAP) was used for data visualization. Trajectory analysis was performed using Slingshot ^28^ (2.0.0) and predicted trajectories were visualised using UMAP. For differential expression analysis between the treated and untreated 48-hour time points, all 4 datasets were normalised using scTransform^29^, 3000 features were selected using the SelectIntegrationFeatures function, and again integrated with the normalization.method parameter set to ‘SCT’.

### Single-cell RNA sequence analysis of irradiated intestinal epithelial cells

Whole small intestines from two irradiated GF or two SPF mice were pooled were flushed with cold PBS and flared longitudinally. Small intestines were sectioned into 1cm pieces and washed with cold PBS two times. Intestinal sections were incubated in 2mM EDTA in PBS at 4°C for 40 mins with continuous shaking. Intestinal sections were washed with cold PBS and crypt enrichment was performed by vigorously shaking intestinal sections in cold PBS and passing the dissociated epithelial segments through a 70-μm cell strainer. Following centrifugation, enriched intestinal crypts were washed with cold PBS and digested with 37°C trypsin (0.25% in HBSS) and mixed by gentle pipetting every 20 minutes for 50 minutes or until single cells were attained. The single cell suspension was washed with ice-cold DMEM and passed through a 40-μm cell strainer. An estimated 10000 cells were loaded for 10x Genomics single-cell isolation and library preparation following manufacturer’s recommendations. Illumina Hiseq 3000 was used to sequence the sample. Sequences were demultiplexed, unique molecular identifiers were aggregated and mapped to the mm10/GRCm38 transcriptome using Cellranger v3.0.0 (10x Genomics). Data generated by Cellranger were used to generate a SingleCellExperiment object. Scater^30^ was used to assess the quality of cells. Briefly, genes expressed in less than five cells were removed. Cells with less than 1000 counts or 600 features, and more than 50% mitochondrial genes were removed. Quality control thresholds were chosen after assessing the impact of each threshold on all cell types. Gene expression data were normalized and integrated using sctransform in Seurat. An elbow plot was used for the selection of principle components. Cells underwent unsupervised clustering using the shared-nearest-neighbour modularity optimization-based clustering algorithm (smart local moving algorithm). The optimal resolution for cell clusters was determined using the clustree package by increasing the clustering resolution by increments of 0.1 from 0 to 1. Cells were visualized using the uniform manifold approximation and projection (UMAP) method. Cell types were annotated using previously defined cell markers ^31^. Immune cells were filtered by expression of *Ptprc*. All scRNAseq analysis was performed using R 3.6.0.

### EdU incorporation and flow cytometry

In brief, ADP-Hep treated organoids were treated with EdU for 2 hours prior to collection. Organoids were dissociated using Trypsin-EDTA (325-043-EL, Wisent) at 37°C for 5 minutes with periodic disruption and suspended cells were passed through a 40μm cell trainer. Samples were stained with the LIVE/DEAD Fixable Aqua Dead Cell Stain kit (L34966, Invitrogen) to label dead cells, followed by the EdU Click-It Flow Cytometry kit (C10419, Invitrogen), as per manufacturer instructions. The samples were analyzed on an LSR Fortessa Cell Analyzer (BD Biosciences) and results were analyzed with FlowJo Version 10.

### Mass Spectrometry

Stool pellets from either SPF or GF mice were pooled and disrupted in extraction solvent (ACN:MeOH:H2O 2:2:1 + 0.1% Formic acid). Insoluble material was removed by centrifugation at 14000*g* at 4°c and supernatants were dried under N2. For analysis samples were reconstituted in 30uL of LCMS grade water. Analytes were separated using an Agilent ZORBAX Extend-C18 reverse phase chromatography using tributylamine as an ion paring agent. Data was analyzed using Agilent’s MassHunter Qualitative Analysis software.

### Statistical Analysis

GraphPad Prism 8 was used for statistical analysis and graph generation. Each point represents a biological replicate. For comparison of two data with 2 groups, unpaired t-tests were performed to determine significance. For comparison of multiple groups unpaired one-way ANOVAs were performed.

---

**Supplemental Figure 1.**
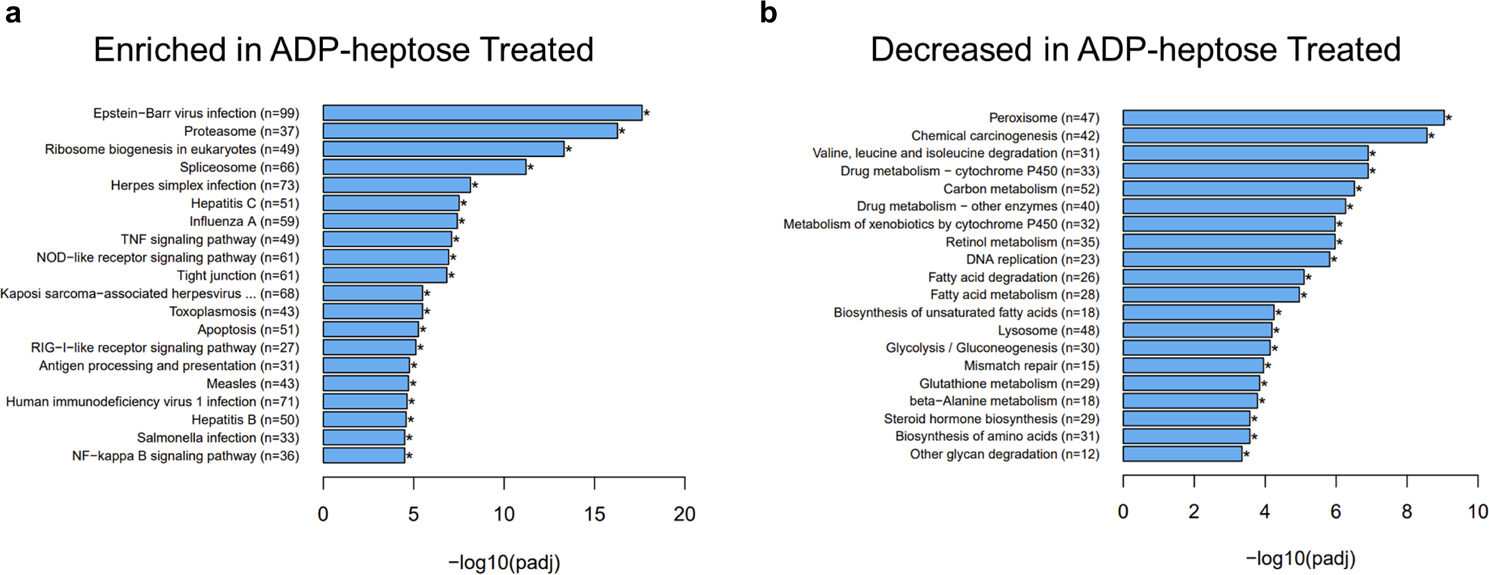
The ALPK1-TIFA axis is expressed and functional within ISCs in the small intestine. **a-b,** Barplot of KEGG enrichment analysis for pathway enriched or decreased in ileal organoids stimulated with ADP-Hep for 6 hours compared to unstimulated organoids.

**Supplemental Figure 2.**
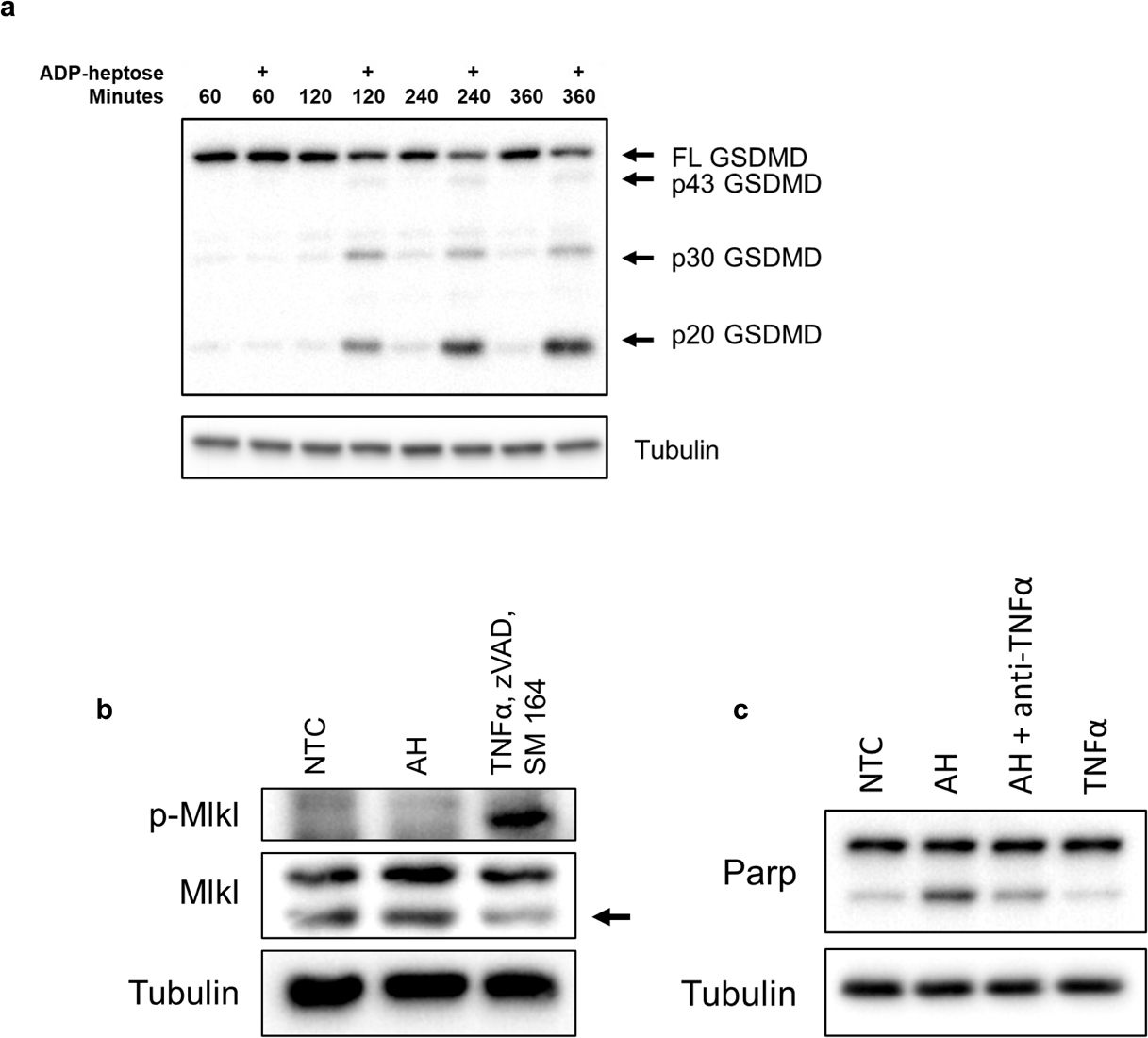
ADP-Hep does not induce pyroptosis or necroptosis. **a,** Immunoblot analysis of ileal organoids treated with ADP-Hep [20µg/mL] for 60, 120, 240 and 360 minutes. **b,** Immunoblot analysis of ileal organoids stimulated for 3 hours with ADP-Hep [20µg/mL] or TNFα [10ng/mL], zVAD [20μM], and SM164 [1 μM]. Arrow indicates Mlkl band. **c,** Immunoblot analysis of ileal organoids treated with ADP-Hep [20µg/mL], or AH [20µg/mL] and anti-TNFα, or TNF [10ng/mL] alone.

**Supplemental Figure 3.**
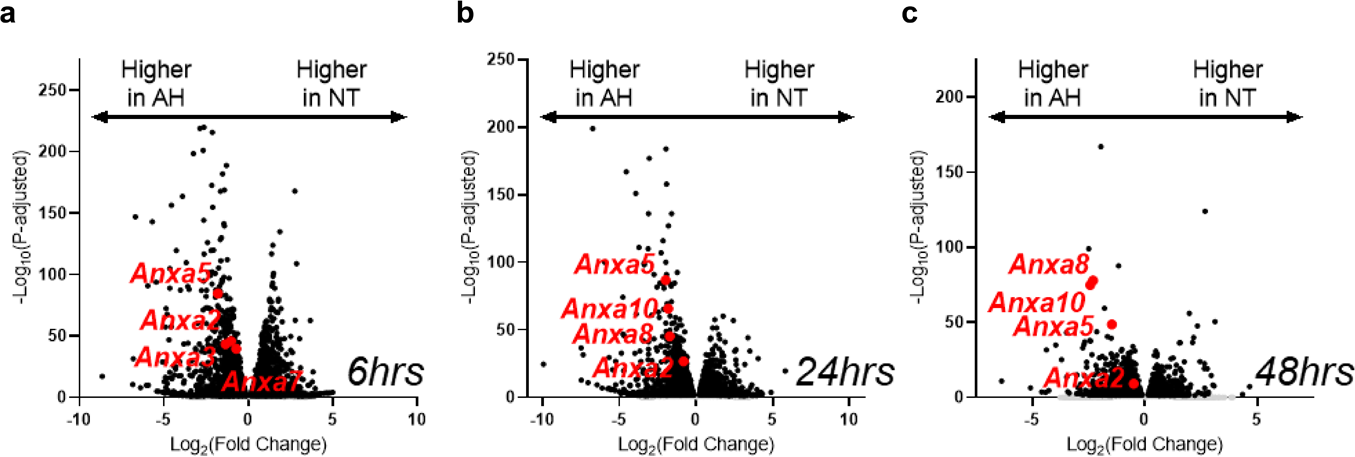
A*n*xa transcripts are induced following ADP-Hep stimulation in ileal organoids. **a-c,** Volcano plot of differential mRNA expression of ileal organoids stimulated with ADP-heptose [20µg/mL] for 6, 24 or 48 hours compared to untreated.

**Supplemental Figure 4.**
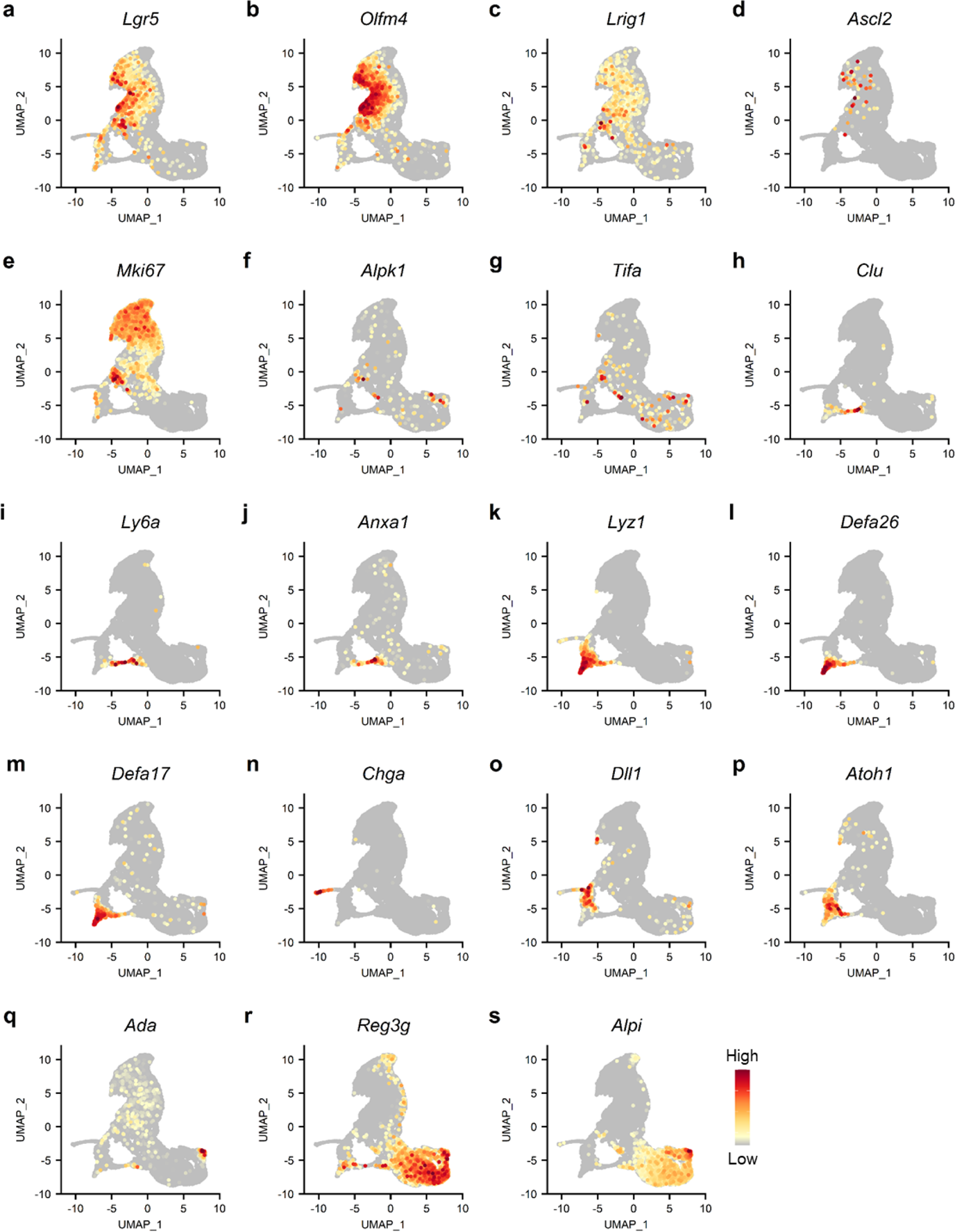
Ileal organoids express conventional intestinal epithelial markers. **a-s,** Expression of indicated genes overlayed on UMAP plots.

**Supplemental Figure 5.**
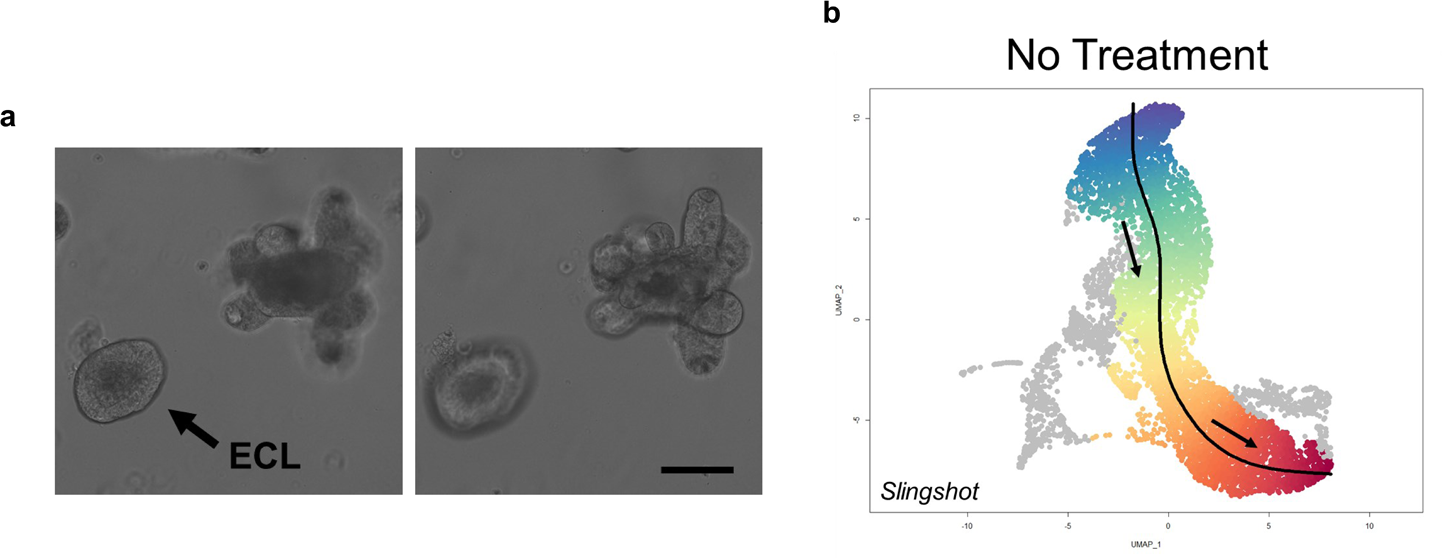
SI organoid cultures contain enterocyst-like organoids. **a,** Brightfield imaging of enterocyst-like organoid present in culture next to mature organoid. **b,** Slingshot analysis showing predicted trajectory of enterocyst-like cells in single cell data set.

**Supplemental Figure 6.**
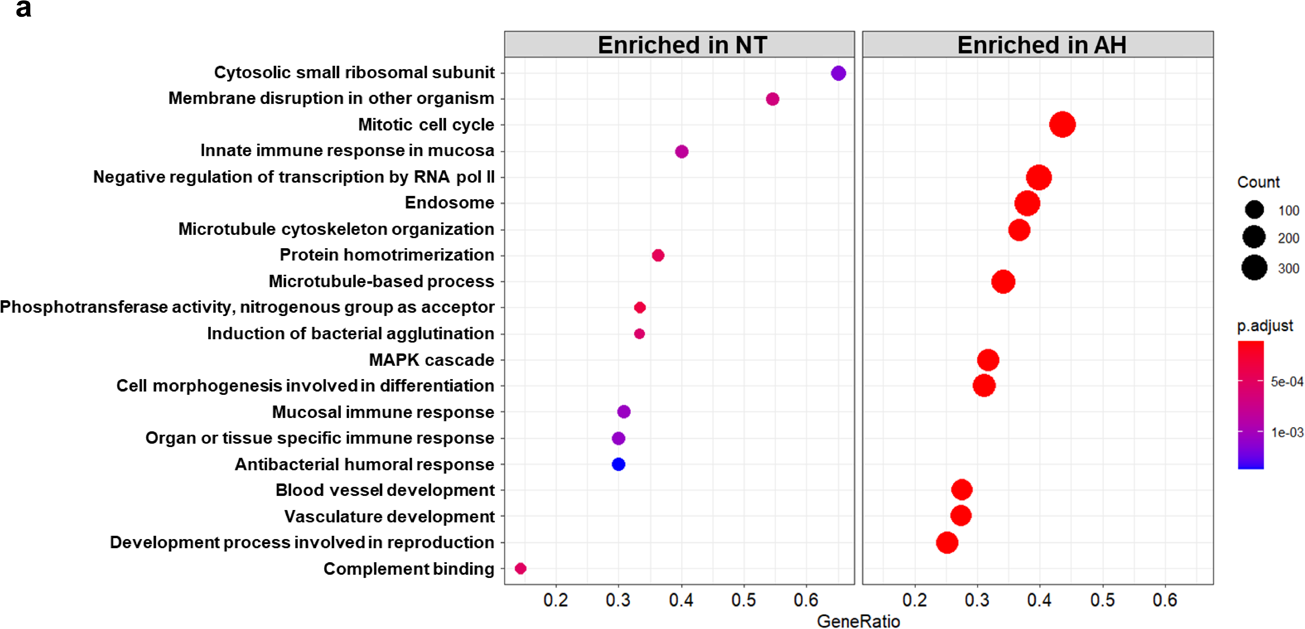
ADP-Hep induces a regenerative program to rederive ISCs from PCs. **a,** GO enrichment analysis of Lyz1^+^ cells from untreated and AH-Hep treated organoids scRNA-seq dataset.

**Supplemental Figure 7.**
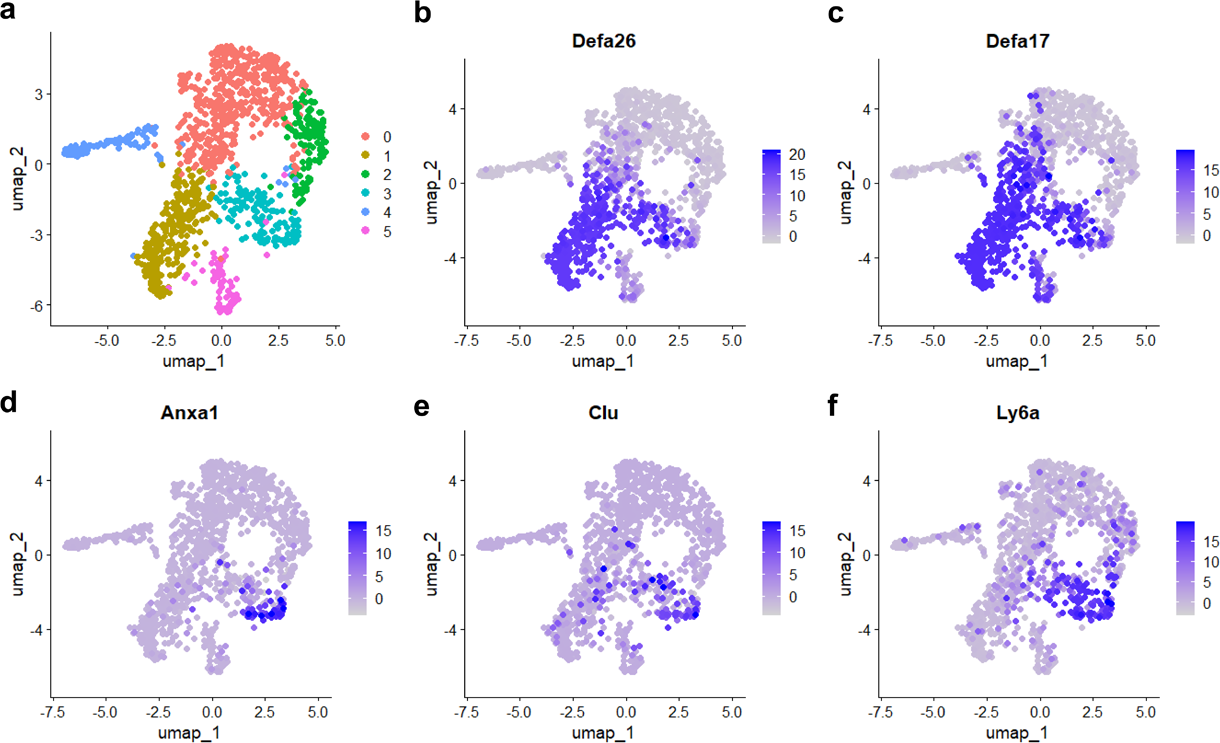
Markers of revSCs are co-expressed with markers of Paneth cell. **a,** UMAP of re-clustered Paneth and goblet cell. **b-f,** Expression of indicated genes overlayed on UMAP plots.

**Supplementary Table 1.**
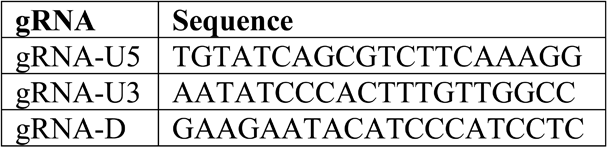
gRNA sequences for Tifa knockout mouse generation.

**Supplementary Table 2.**
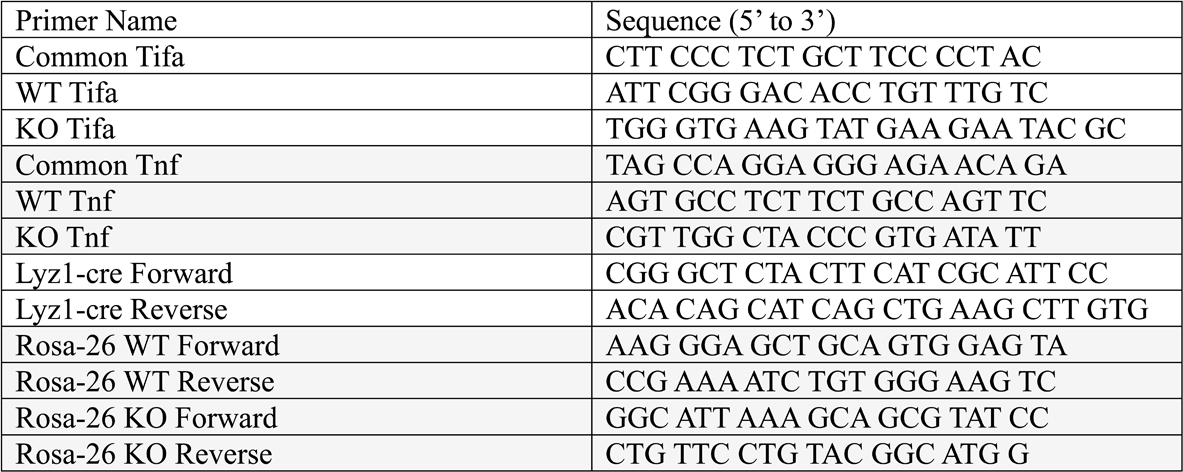
Genotyping Primers.

**Supplementary Table 3.**
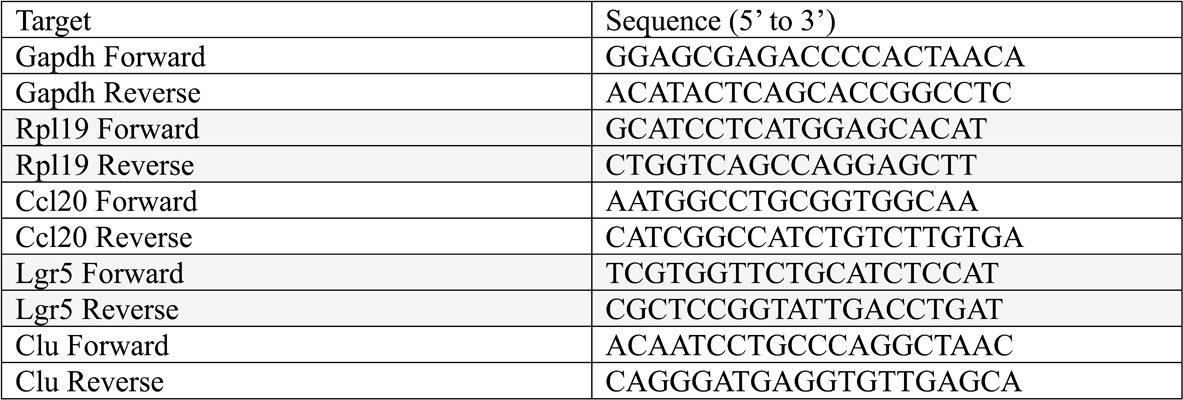
Murine qPCR primers used in this study.

